# Increased mRNA expression of CDKN2A is a transcriptomic marker of clinically aggressive meningiomas

**DOI:** 10.1101/2022.12.07.519545

**Authors:** Justin Z. Wang, Vikas Patil, Jeff Liu, Helin Dogin, Felix Behling, Marco Skardelly, Marcos Tatagiba, Elgin Hoffman, Severa Bunda, Rebecca Yakubov, Ramneet Kaloti, Sebastian Brandner, Andrew Gao, Aaron-Cohen Gadol, Jennifer Barnholtz-Sloan, David Raleigh, Felix Sahm, Paul C. Boutros, Ghazaleh Tabatabai, Kenneth Aldape, Farshad Nassiri, Gelareh Zadeh

## Abstract

**Background:** Homozygous loss of CDKN2A/B is a genetic alteration found in many cancer types including meningiomas, where it is associated with poor clinical outcome. It is now also a diagnostic criterion for grade 3 meningiomas in the 2021 WHO classification for central nervous system tumors. However, as in other cancers, the relationship between copy number loss of CDKN2A/B and expression of its gene product is unclear and may be either commensurate or paradoxical in nature. Therefore, we aimed to investigate the association of CDKN2A mRNA expression with clinical prognosis, WHO grade, and other molecular biomarkers in meningiomas such as DNA methylation, molecular group, and proteomics.

**Methods:** We used multidimensional molecular data of 490 meningioma samples from 4 independent cohorts to examine the relationship between mRNA expression of CDKN2A and copy number status, its correlation to clinical outcome, the transcriptomic pathways altered in differential CDKN2A expression, and its relationship with DNA methylation, and proteomics using an integrated molecular approach.

**Results:** Meningiomas without any copy number loss were dichotomized into high (CDKN2A^high^) and low (CDKN2A^low^) CDKN2A mRNA expression groups. Patients with CDKN2A^high^ meningiomas had poorer progression free survival (PFS) compared to those with CDKN2A^low^ meningiomas. CDKN2A mRNA expression was increased in more aggressive molecular groups, and in higher WHO grade meningiomas across all cohorts. CDKN2A^high^ meningiomas and meningiomas with CDKN2A copy number loss shared common up-regulated cell cycling pathways. CDK4 mRNA expression was increased in CDKN2A^high^ meningiomas and both p16 and CDK4 protein were more abundant in CDKN2A^high^ meningiomas. CDKN2A^high^ meningiomas were frequently hypermethylated at the gene body and UTR compared to CDKN2A^low^ meningiomas and found be more commonly Rb-deficient.

**Conclusions:** An intermediate level of CDKN2A mRNA expression appears to be optimal as significantly low (CDKN2A deleted) or high expression (CDKN2A^high^) are associated with poorer outcomes clinically. Though CDK4 is elevated in CDKN2A^high^ meningiomas, Rb-deficiency may be more common in this group, leading to lack of response to CDK inhibitors.

## Introduction

Meningiomas are the most common primary intracranial tumour in adults.^1^ Though the majority of meningiomas are benign, 20-30% are clinically aggressive and have a tendency to recur despite maximal standard of care therapy with surgical resection and radiotherapy.^2-5^ Compared to benign meningiomas, aggressive meningiomas tend to harbour a greater burden of copy number changes including focal deletions.^2,6^ In particular, homozygous deletion of the cyclin-dependent kinase inhibitor 2A/B (CDKN2A/B) genes on chromosome 9p21 have been demonstrated to be enriched in higher WHO grade or recurrent meningiomas, and when present, has been associated with significantly shorter time to progression compared to tumours without this deletion.^7-10^ However, even in cohorts enriched for WHO grades 2 and 3 meningiomas, homozygous CDKN2A/B deletion is rare, reported in only 1.7-4.9% of patients.^2,8,11^ Despite the relative rarity of this finding, its predictive value for poor prognosis has led to its incorporation into the 2021 central nervous system (CNS) WHO classification as a diagnostic criterium for grade 3 meningiomas.^12^

Although copy number loss of CDKN2A/B has been implicated in many different cancers, including meningiomas, how this copy number change correlates with mRNA expression, and the prognostic significance of this mRNA expression has at times yielded paradoxical findings.^13-17^ For example, whereas decreased mRNA expression of CDKN2A (p16) has been found to be a poor prognostic factor in high-grade gliomas, increased expression has been associated with worse outcomes in other cancers such as ovarian and bladder cancer.^13,18,19^ Whether these seemingly discordant relationships are cancer/tissue-specific or due to common alterations independent of- or downstream to the p16 pathway remain to be determined.^20^

Recent work from our group integrated multiple epigenomic and genomic platforms to uncover four consensus molecular groups (MG) of meningiomas, each with unique biology and outcomes.^3^ We found that several meningiomas without CDKN2A copy number loss had low levels of CDKN2A mRNA expression, comparable to meningiomas with CDKN2A loss, yet had much better clinical outcomes. This prompted us to examine the relationship between CDKN2A mRNA expression and copy number status, WHO grade, MG, transcriptomic pathways, DNA methylation, proteomics, and clinical outcome using our published data as the discovery cohort, and validating these findings using a novel independent cohort from Tübingen, and two large publicly available cohorts from recently published studies.^21,22^

## Methods

### Patient samples and clinical annotation

491 clinically annotated meningiomas with matched molecular data were used in this study **(Supplementary Table 1)**. We used 122 tumors from Toronto (Canada) with previously published multiplatform genomic and epigenomic data available as the discovery cohort, 75 new samples from Tübingen (Germany), enriched for clinically aggressive meningiomas, and publicly available cohorts from recently published studies by Bayley et al. and Choudhury et al. comprised of 109 and 185 meningiomas as the validation cohorts.^3,6^ Available clinical data were collected based on previously determined consensus core data elements for meningiomas in the Tübingen cohort.^23^ Each case was reviewed centrally by two independent neuropathologists to confirm the diagnosis of a meningioma and was graded based on the 2016 WHO classification criteria. Tumour recurrence and extent of resection were defined consistent with our previous work.^3^ Clinical data for the Bayley et al. and Choudhury et al. studies were obtained where publicly available as Supplementary Materials.^21,22,24^

### DNA and RNA extraction

DNA and RNA were extracted from meningiomas of the Tübingen validation cohort as detailed previously using the DNeasy Blood and Tissue Kit (Qiagen, USA) and RNeasy Mini Kit (Qiagen, USA) as previously described.^3^ Approximately 250-500 ng of extracted DNA as quantified on the Nanodrop 1000 Instrument (Thermo Scientific, USA) were bisulfite converted (EZ DNA Methylation Kit, Zymo, California, USA). RNA with RNA Integrity Number (RIN) > 7 when assessed using the Agilent 2100 Bioanalyzer (Agilent, USA) were selected to move forward for further sequencing.

### DNA methylation

DNA methylation data were generated as previously described using the Illumina Infinium MethylationEPIC BeadChip Array (Illumina, San Diego, USA).^3^ Data processing was conducted as previously published.^3^ Briefly, raw data files (*.idat) were imported, processed, and normalized. General quality control measures were performed as per the manufacturer instructions. Methylation probe annotation was performed using the University of California Santa Cruz Genome Browser (GRCh38/hg38 assembly). Copy number alterations were inferred from the methylation data using conumee.^25^ A log2 value of -0.4 was utilized as a cut-off below which we determined a homozygous deletion/loss (homodel) at the CDKN2A/B locus.^26^ A log2 value between -0.2 and -0.4 was utilized to call a segmental or partial deletion/loss (partialdel) at that specific locus. Beta-value at CpG sites annotated to the specific genes including their gene body, UTR, and promoter were obtained after filtering out probes that failed to pass our mean detection threshold.

### Whole exome sequencing (WES)

Exome data preparation, quality assessment, and analysis were as previously published.^3^ Raw sequencing data (fastq) were aligned to the hg19 reference genome using BWA-MEM v0.7.12.^27^ Sequenza v2.1.2 were used for tumour exome samples with matching blood while CNVkit v0.9.6 were used for tumour samples without matched blood using a pooled reference set of 60 peripheral blood samples from individuals not included this study.^28,29^ We utilized the previously published cut-off log_2_ratio <-0.7 as a homodel (or deep deletion) at the CDKN2A/B locus, while a log_2_ratio from -0.35 to -0.7 was categorized as a heterozygous deletion (heterodel).^30^

### RNA sequencing

mRNA libraries were generated as previously described using the NEB Ultra II directional mRNA library prep kit in accordance with the manufacturer’s instructions.^3^ Libraries were sequenced on the Illumina HiSeq 2500 high output flow cell (2×126bp), sequenced with 3 samples per lane to obtain ∼70 million reads per sample. Raw sequencing files (fastq) were processed and aligned to the human reference genome GRCh38 using STAR (v2.6.0a).^31^ Duplicate reads were removed using SamTools (v1.3).^32^ Raw gene expression counts were calculated for each sample using Rsubread (v1.5.0), normalized by counts-per-million, and subjected to trimmed mean of M normalization using edgeR (v3.22.3).^33,34^ CPM values were converted to Z-scores for each sample for subsequent analysis. Samples with CDKN2A expression Z-score ≥ 1 in each cohort were designated as meningiomas with high CDKN2A expression (CDKN2A^high^), while the remaining were designated to have low expression (CDKN2A^low^). Differential RNA sequencing analysis was conducted using limma with multiple false discovery rate (FDR) and adjusted p-value cut-offs where indicated.^35^

### RNA pathway analysis

Pathway enrichment analysis and visualization of pathway data were performed as previously described.^36^ Pathway enrichment analysis were defined by the pathway gene sets from http://baderlab.org/GeneSets, which are updated monthly and performed as previously published.^3,36^ Results of pathway enrichment analysis were visualized using Cytoscape (v3.7.2).^37^ Network maps were generated for nodes with p-value <0.0001. Nodes sharing overlapping genes (Jaccard Coefficient >0.25) were connected with a green edge. Pathways were grouped together based on shared keywords in description of the pathways using AutoAnnotate (v1.2) and manually through mechanistic similarities if they did not fit into a specific pathway group automatically.^36^

### Proteomics

Protein data were generated through shotgun proteomics as previously described using an Orbitrap Fusion (Thermo Scientific) tribid mass spectrometer.^3^ Peptides were detected using a Top25 data-dependent acquisition method. Data were reviewed against a UniProt complete human protein sequence database with an FDR of 1% for peptide spectral matches. Relative label-free protein quantitation was calculated using MS^1^-level peak integration and with matching-between-runs feature. Proteins identified with a minimum of two peptides were used for subsequent analysis.

### pRB Western Blot

For western blotting, tissue samples were homogenized and lysed in EBC buffer (50 mM Tris [pH 8.0], 120 mM NaCl, 0.5% Nonidet P [NP]-40) supplemented with protease and phosphatase inhibitors. Proteins were eluted by boiling in sample buffer and resolved by SDS–polyacrylamide gel electrophoresis. Proteins were electro-transferred onto polyvinylidene difluoride membrane (Bio-Rad), blocked and probed with the indicated antibodies (Rb Antibody Sampler Kit #9969) and β-actin from Cell Signaling Technologies (Danvers, Massachusetts, USA).

### p16 Immunohistochemistry

Immunohistochemistry (IHC) was carried out on 5 μm paraffin sections of a separate independent cohort of meningiomas (DKFZ, Heidelberg, Germany) from mounted on poly-L-lysine coated slides. Antibody against p16 (DAKO, 1:25, positive control human tonsil tissue) were utilized with standard techniques. All slides were counterstained with hematoxylin.

### Publicly available datasets

We downloaded raw methylation and RNA-seq data from the Gene Expression Omnibus database https://ncbi.nlm.nih.gov/geo, under the following accession numbers: GSE189521 (Bayley et al. DNA methylation), GSE189672 (Bayley et al. RNAseq), GSE183656 (Choudhury et al. DNA methylation and RNAseq), and accompanying clinical data sample sheets from the aforementioned manuscripts.^21,22^

### Molecular Group designation

Consensus MG designation as described above were obtained from integration of DNA methylation, RNA expression, and copy number alteration data on the Toronto cohort of our previous study.^3^ MG designation in the Tübingen, Bayley et al. and Choudhury cohorts were determined using the Metagene model building trained on the unique transcriptional signatures of each MG from the Toronto discovery cohort to probabilistically predict the most likely MG for each tumour.^38^ Methylation group prediction in the Bayley et al. (MenG A, B, C) and Choudhury et al. (MethG Merlin-Intact (MI), Immune-enriched (IE), Hypermitotic (HM)) cohorts, using on their own respective groups as the reference for one another, and for the Toronto and Tübingen cohorts were performed in a similar manner using a Metagene model trained on the unique CpGs of each methylation group from their original publication to predict the methylation group of the other cohorts **(Supplementary materials)**.

### Statistics

Chi-square test of proportions were used to compare proportions between two groups. Continuous clinical variables between two groups were compared using Welch’s t-test. Comparison of RNA expression or protein abundance between two groups were done using the Wilcoxon-Mann-Whitney U test. Comparison of continuous variables between multiple groups were performed using the Kruskal-Wallis one-way analysis of variance (ANOVA), followed by post-hoc Dunn’s test. For survival analysis, Kaplan-Meier (KM) survival plots were generated using the package survminer and log-rank tests were done to test the null hypothesis that there were no difference in progression free survival (PFS) between groups. Univariable and multivariable survival analyses were conducted by fitting a Cox proportional hazards models with the clinical covariates on the combined clinical cohort. The proportional hazards assumption was tested using the *ggcoxzph* function in the *survminer* R package and by plotting the scaled Schoenfeld residuals of each covariate against transformed time. Hazard ratios and 95% confidence intervals were reported. Pearson correlation test was used to test the correlation between two variables with reporting of the correlation coefficient and p-value. Statistical significance for all tests were set at p<0.05 unless otherwise specified.

## Results

### Clinical cohort

The Toronto discovery cohort included 122 meningiomas with tumour DNA methylation, 121 of which had matched WES, RNA sequencing (RNAseq) data, and 96 with bulk proteomics data.^3^ The Tübingen validation cohort consisted of 75 meningiomas with RNAseq, 60 of which had matched DNA methylation data. The publicly available datasets were comprised of 109 meningiomas with matched DNA methylation and RNAseq in the Bayley et al. cohort and 185 meningiomas in the Choudhury et al. cohort with the same matched datasets **(Supplementary table 1)**.^21,22^ Baseline characteristics of these cohorts are summarized in **Table 1**.

**Table 1.** Patient characteristics of the Toronto discovery cohort, Tübingen validation cohort, and publicly available cohorts (Bayley et al., Choudhury et al.) stratified by CDKN2A expression.

|  | Toronto Discovery Cohort (n=122) |  | Tübingen Validation Cohort (n=75) |  | Bayley et al. Cohort (n=109) |  | Choudhury et al. Cohort (n=185) |  |
| --- | --- | --- | --- | --- | --- | --- | --- | --- |
| CDKN2A Copy Number Status |  |  |  |  |  |  |  |  |
| Homodel | 6 (5%) |  | 0 |  | 1 (1%) |  | 11 (6%) |  |
| Heterodel/Partialdel | 6 (5%) |  | 1 (1%) |  | 1 (1%) |  | 5 (3%) |  |
| Nondel | 110 (90%) |  | 74 (99%) |  | 107 (98%) |  | 169 (91%) |  |
| CDKN2A Expression Group |  |  |  |  |  |  |  |  |
|  | Low (n=91) | High (n=19) | Low (n=64) | High (n=10) | Low (n=90) | High (n=17) | Low (n=137) | High (n=32) |
| Gender |  |  |  |  |  |  |  |  |
| Male | 37 (41%) | 7 (37%) | 26 (41%) | 3 (30%) | 32 (36%) | 8 (47%) | 46 (34%) | 10 (31%) |
| Female | 54 (59%) | 12 (63%) | 38 (59%) | 7 (70%) | 58 (64%) | 9 (53%) | 91 (66%) | 22 (69%) |
| WHO grade |  |  |  |  |  |  |  |  |
| 1 | 51 (56%) | 4 (21%) | 20 (31%) | 3 (30%) | 77 (86%) | 12 (71%) | 76 (55%) | 9 (28%) |
| 2 | 28 (31%) | 10 (53%) | 42 (65%) | 7 (70%) | 13 (14%) | 5 (29%) | 53 (39%) | 15 (47%) |
| 3 | 12 (13%) | 5 (26%) | 2 (4%) | 0 | 0 | 0 | 8 (6%) | 8 (25%) |

| **Extent of Resection** |  |  |  |  |  |  |  |  |
| --- | --- | --- | --- | --- | --- | --- | --- | --- |
| GTR | 65 (72%) | 11 (58%) | 37 (58%) | 5 (50%) | 69 (77%) | 13 (76%) | NA | NA |
| STR | 26 (28%) | 8 (42%) | 27 (42%) | 5 (50%) | 21 (23%) | 3 (18%) | NA | NA |
| **Molecular Group** |  |  |  |  |  |  |  |  |
| MG1 | 14 (15%) | 3 (16%) | 5 (8%) | 0 |  |  |  |  |
| MG2 | 28 (30%) | 0 | 25 (39%) | 1 (10%) |  |  |  |  |
| MG3 | 36 (40%) | 5 (26%) | 25 (39%) | 2 (20%) |  |  |  |  |
| MG4 | 13 (15%) | 11 (58%) | 9 (14%) | 7 (70%) |  |  |  |  |
| MenG A |  |  |  |  | 47 (52%) | 4 (23%) |  |  |
| MenG B |  |  |  |  | 23 (26%) | 3 (18%) |  |  |
| MenG C |  |  |  |  | 13 (14%) | 7 (41%) |  |  |
| MI |  |  |  |  |  |  | 63 (46%) | 8 (25%) |
| IE |  |  |  |  |  |  | 47 (34%) | 7 (22%) |
| HM |  |  |  |  |  |  | 27 (20%) | 17 (53%) |

\* Statistical significance with $p < 0.05$ comparing low- versus high-CDKN2A expression groups on 2-sample $\chi^2$ test for equality of proportions with continuity correction for categorical variables or Welch two-sample t-test for continuous variables

### Loss of CDKN2A/B is a rare event in meningiomas and is associated with poor outcome

Loss of CDKN2A/B (homodel, heterodel, or partialdel) was a rare event across all cohorts. Using copy number calling based on WES and DNA methylation in the Toronto discovery cohort, only 6 patients had homodel of CDKN2A/B (4.9%) while 6 (4.9%) had heterodel/partialdel. Copy number calling was performed using DNA methylation for the other cohorts. In the Tübingen cohort, no meningiomas had CDKN2A/B homodel and only 1 had CDKN2A/B partialdel (1.3%), despite this cohort being enriched for highly aggressive meningiomas with early recurrences. In the Bayley et al. cohort 1 meningioma had CDKN2A partialdel (0.9%) and 1 (0.9%) had homodel. In the Choudhury et al. cohort, 5 meningiomas had CDKN2A partialdel (2.7%), while 11 had homodel (5.9%).

While CDKN2A/B homodel is now a diagnostic criterion for WHO grade 3 meningiomas, of the meningiomas with this alteration, only 10/18 (56%) were initially classified as WHO grade 3 (Toronto n=2, Choudhury et al. n=8). The majority of WHO grade 3 meningiomas did not have any CDKN2A loss (35/47, 74%). However nearly all meningiomas with any CDKN2A deletion (homodel/heterodel/partialdel) had poor clinical outcomes **(Supplementary figure 1a**,**b)**. There was no significant difference in outcome between meningiomas with CDKN2A/B homodel compared to those with heterodel in the Toronto cohort (median PFS 1.61 vs 2.00 years, pairwise Log-rank test, adj. p = 0.52, **Supplementary figure 1a)** or partialdel in the combined validation cohorts (median PFS 0.95 vs 2.12 years, pairwise Log-rank test, adj. p = 0.674, **Supplementary figure 1b)**.

### CDKN2A mRNA expression is decreased with homozygous loss

As CDKN2A and CDKN2B mRNA expression were found to be highly correlated with one another in both the Toronto discovery cohort and the combined validation cohorts (Pearson correlation R=0.79, p<2.2×10^−6^; R=0.68, p<2.2×10^−6^, **Supplementary figure 1b**,**c)**, we focused our subsequent analysis on CDKN2A expression. Meningiomas with CDKN2A/B homodel had significantly decreased CDKN2A mRNA expression compared to those without any loss in both the discovery and combined validation cohorts **(Supplementary figure 1d**,**e)**. Interestingly, although meningiomas with CDKN2A heterodel/partialdel had similarly poor clinical outcomes as those with CDKN2A homodel, their mRNA expression appeared to be more heterogeneous and not significantly different from meningiomas without any CDKN2A loss **(Supplementary Figure 1a**,**b**,**d**,**e)**.

### High CDKN2A expression results in poorer outcomes independent of copy number loss

In each cohort, meningiomas were dichotomized into 2 transcriptomic groups (see Methods) with different levels of CDKN2A mRNA expression (hereafter referred to as CDKN2A^high^ and CDKN2A^low^). To establish the prognostic role of CDKN2A mRNA expression independent of CDKN2A deletions, meningiomas with any copy number loss of CDKN2A (homodel, heterodel, or partialdel) were excluded from these 2 transcriptomic groups. Pooled mRNA expression counts across all cohorts demonstrated that CDKN2A^low^ meningiomas had an intermediate level of CDKN2A expression between those with CDKN2A homodel and CDKN2A^high^ **(Supplementary figure 3a)**. CDKN2A^low^ meningiomas also had the longest PFS, superior to meningiomas with CDKN2A deletions and CDKN2A^high^ meningiomas **(Figure 1a-e)**. This relationship also held true within each WHO grade **(Figure 1g)**.

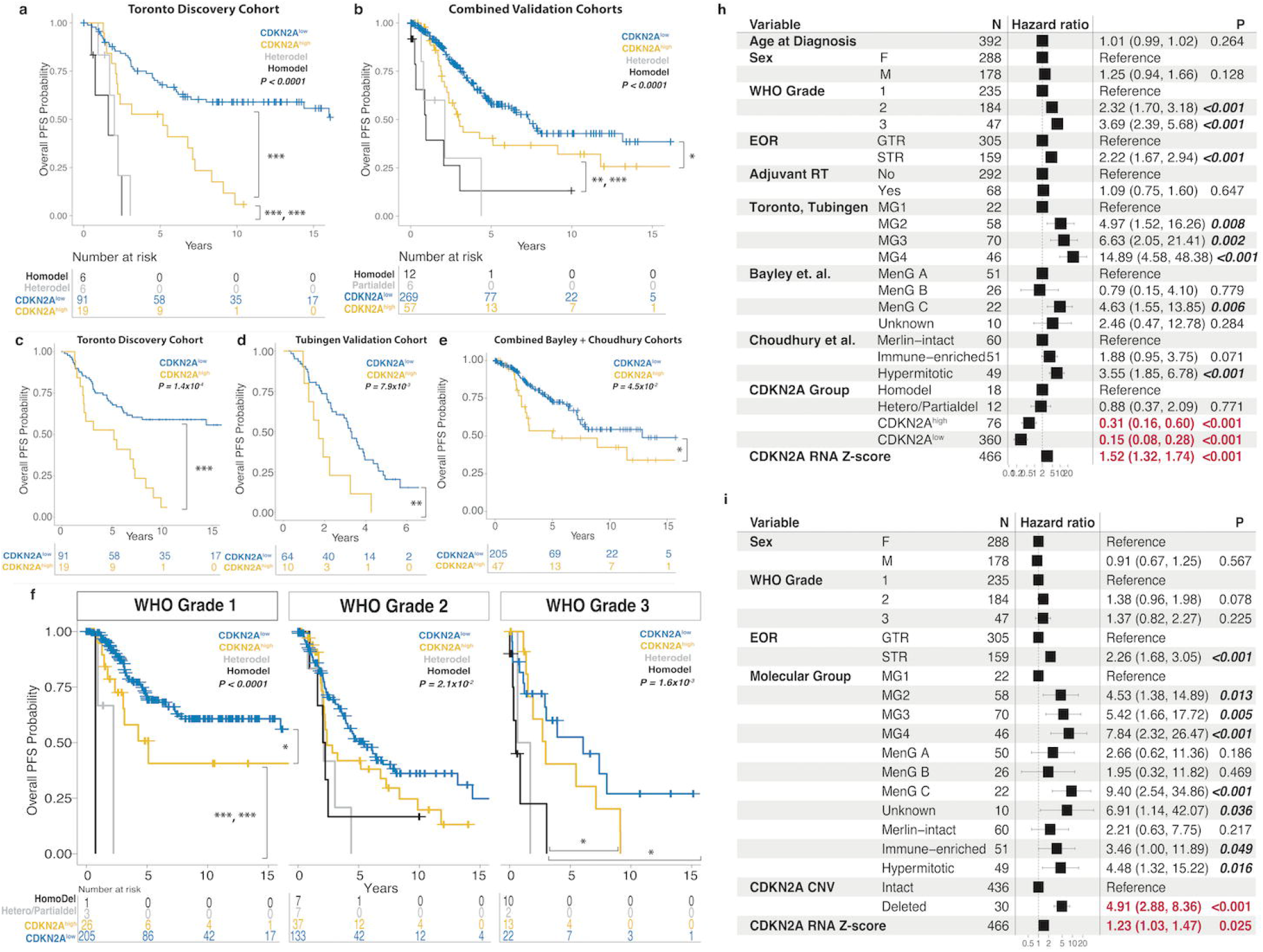

To confirm that CDKN2A mRNA expression could be predictive of outcome, we first fit a univariable **(Figure 1h)** Cox proportional hazards model on all meningiomas for which PFS data were available. In this univariable analysis, covariates associated with worse PFS included higher WHO grade (grades 2, 3 vs grade 1), subtotal resection (STR) (vs gross total resection; GTR), and more aggressive molecular group (MG2-4 vs MG1; MenG C vs MenG A; Hypermitotic (HM) vs Merlin-intact(MI)) **(Figure 1h)**. In keeping with the KM analysis, CDKN2A^high^ (HR 0.31, 95% CI 0.16-0.60, p=5.48×10^−4^) and CDKN2A^low^ (HR 0.15, 95% CI 0.08-0.28, p=6.10×10^−9^) meningiomas were both associated with significantly better PFS compared to meningiomas with CDKN2A homodel, with CDKN2A^low^ meningiomas having the most favourable outcomes **(Figure 1h)**.

To determine if increased CDKN2A mRNA expression as a continuous variable, independent of cohort or transcriptomic group could be predictive of outcome, we included CDKN2A expression (Z-score) as a covariate in both the univariable and multivariable Cox proportional hazards models **(Supplementary figure 2)**. CDKN2A mRNA Z-score appeared to be significantly associated with worse PFS, including on multivariable analysis where CDKN2A deletion status, and other clinical factors, was controlled for (HR 1.23, 95% CI 1.03-1.47, p = 0.025, **Figure 1i)**. Other covariates significantly associated with worse PFS on multivariable analysis included STR (vs GTR), CDKN2A deletion (homodel, heterodel, or partialdel), and having a more aggressive molecular/methylation group designation (MG2-4, MenG C, HM, or IE vs MG1).

### CDKN2A expression increases with biological aggressiveness and increasing WHO grade

Given the association of increased CDKN2A mRNA expression with poorer outcome, we wanted to ascertain whether there would be differences in expression with more aggressive MG/MenG/MethG and WHO grade.^3,21,22^ When compared to the other MG in the Toronto cohort, MG4 meningiomas had the highest proportion of both CDKN2A^high^ (11/19, 58%) and CDKN2A homodel meningiomas (3/5, 60%, **Figure 2a, left)**. When we treated CDKN2A mRNA expression as a continuous variable, we found higher CDKN2A expression in the MGs with the most biologically aggressive meningiomas (MG3 and MG4), with MG4 meningiomas having the highest overall level of expression (**Figure 2a, right**). These findings were corroborated in the Tübingen cohort **(Figure 2b)**. In the Bayley et al. cohort, their most aggressive MenG C contained the only meningiomas with CDKN2A deletions in their cohort, the highest proportion of CDKN2A^high^ meningiomas (n=7) and had the highest level of CDKN2A expression **(Figure 2c)**. In the Choudhury et al. cohort, their HM group, also comprised of the most treatment-refractory meningiomas, had the highest proportion of CDKN2A homodel (n=6), CDKN2A^high^ meningiomas (n=17), and the highest CDKN2A mRNA expression **(Figure 2d)**. These results were reproduced even when all samples were combined and grouped based on each study’s respective molecular classification using a modeling approach (See Methods, **Figure 2e-g)**.

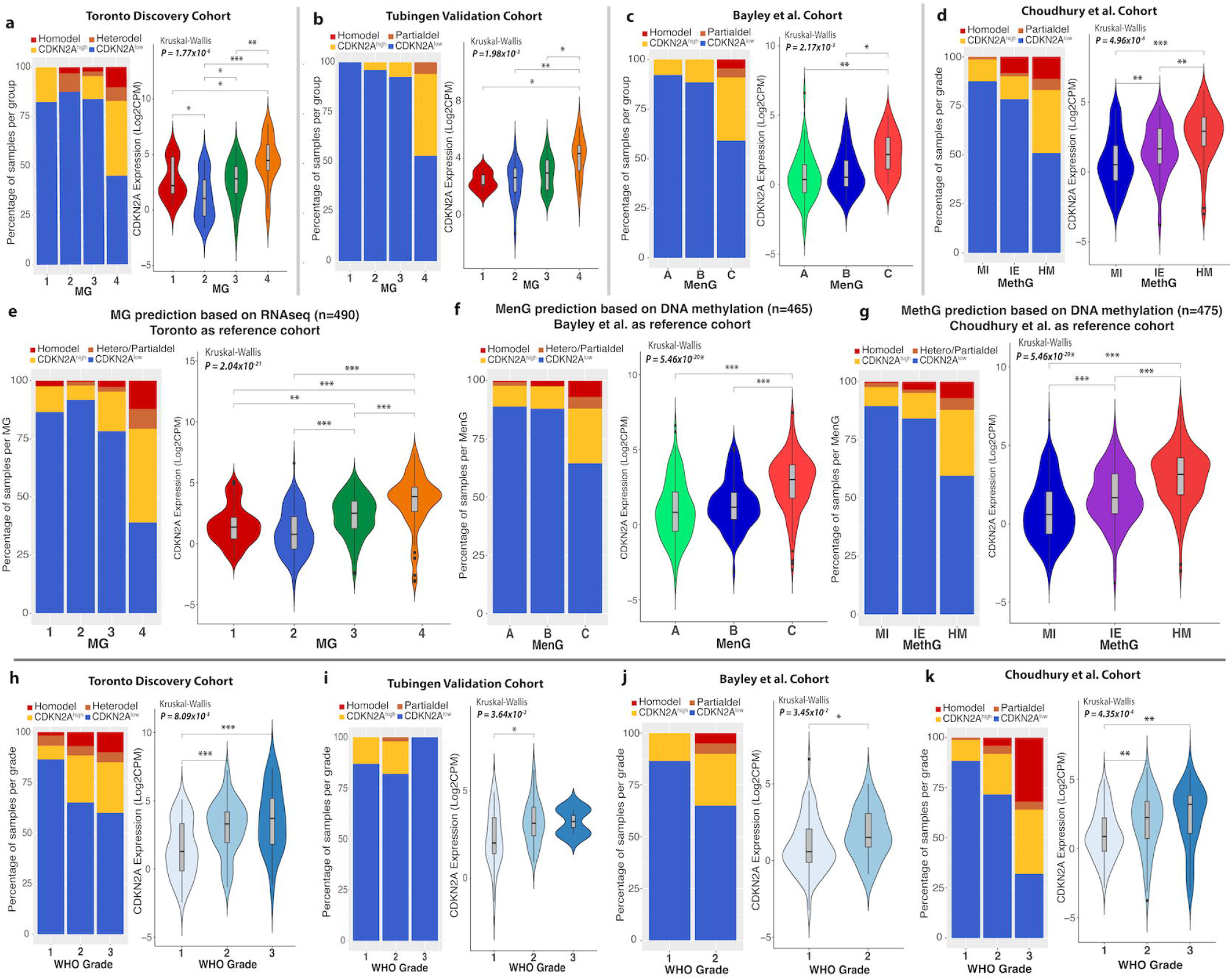

When we stratified meningiomas by WHO grade, we found a similar trend of increasing CDKN2A mRNA expression with higher WHO grade, with WHO grade 2 meningiomas showing significantly higher CDKN2A expression compared to grade 1 tumours across all 4 cohorts **(Figure 2h-k)**. WHO grade 3 meningiomas also appeared to have the highest levels of CDKN2A expression in the Toronto and Choudhury et al. cohorts **(Figure 2h**,**k)**. As there were only 2 WHO grade 3 meningiomas in the Tübingen cohort and none in the Bayley et al. cohort, meaningful comparisons with grade 3 tumours could not be made in these cohorts.

These findings, together with the above however, suggest that regardless of stratification by molecular classification or WHO grade, CDKN2A mRNA expression is elevated in more biologically aggressive meningiomas.

### Multiple transcriptomic pathways involved in both cell cycle control and progression are upregulated in CDKN2A^high^ meningiomas

To explore the transcriptomic differences between CDKN2A^high^ and CDKN2A^low^ meningiomas, we performed differential RNAseq analysis between these groups in each cohort. There was high concordance in the top significantly up- and down-regulated genes observed in at least 2 cohorts **(Figure 3a)**. These genes corresponded to common upregulated pathways involved in cell cycle progression and control at the G1-S transition in CDKN2A^high^ meningiomas compared to their CDKN2A^low^ counterparts across all 4 cohorts **(Figure 3b)**. Pathway enrichment analysis showed convergence of upregulated transcriptomic pathways involved in mitoses, cell cycling, cell cycle control, and apoptosis in CDKN2A^high^ meningiomas across cohorts and downregulation of endothelial, vascular, and metabolic-related pathways **(Figure 3d, e)**.

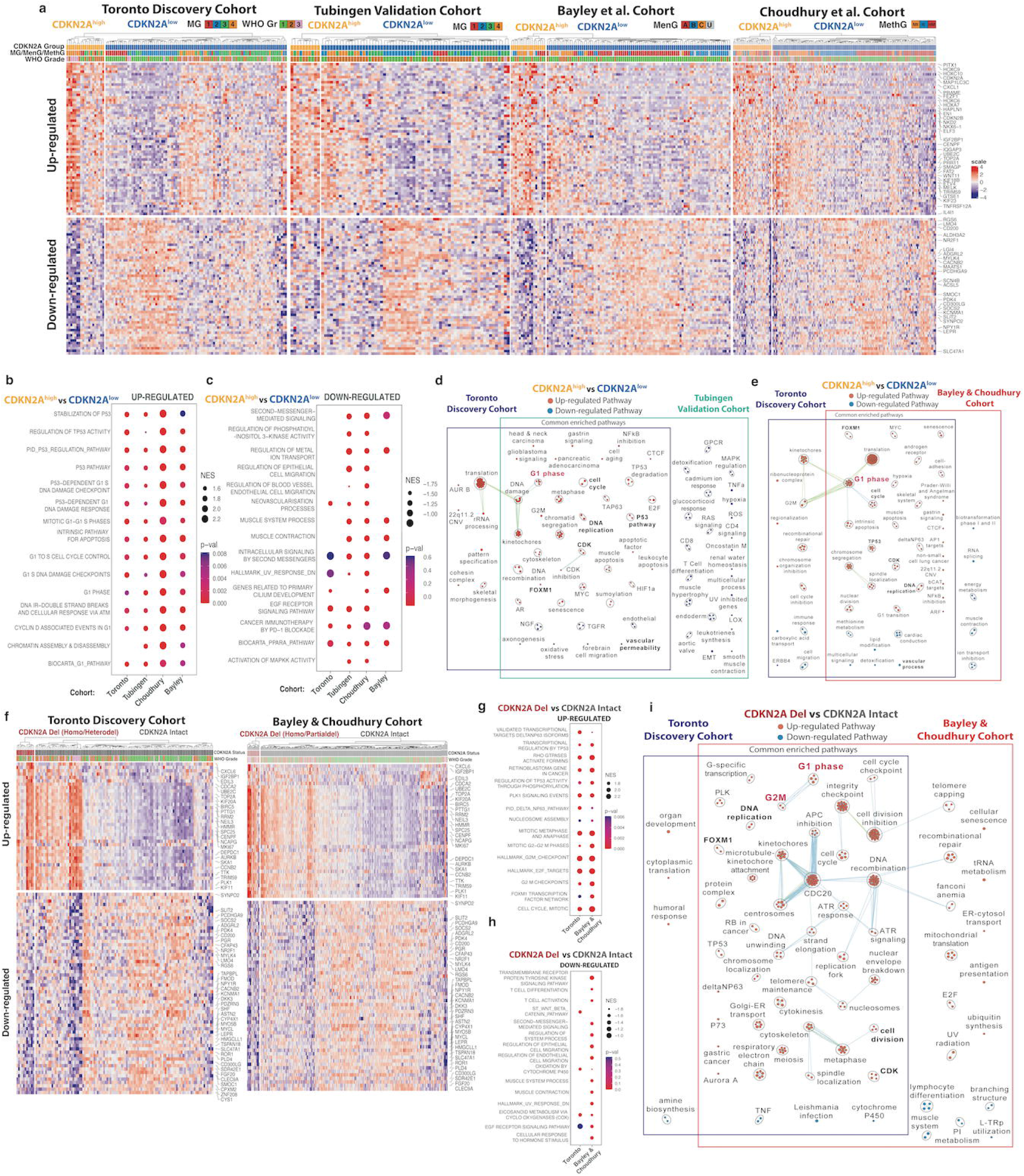

### CDKN2A^high^ meningiomas share transcriptomic pathways with meningiomas that have copy number loss of CDKN2A

CDKN2A^high^ meningiomas and meningiomas with focal CDKN2A deletions (homodel, heterodel, or partialdel) were mutually exclusive in our study and despite the discordant status of the CDKN2A gene, they both shared poor clinical outcomes. Therefore, we wanted to explore whether similar transcriptomic pathways may underlie this shared biological aggressiveness. The Bayley et al. cohort was combined with the Choudhury et al. cohort for this analysis due to the relative scarcity of meningiomas with CDKN2A copy number loss in the former study (n=2). We saw significant up-regulation of similar cell cycle pathways in meningiomas with CDKN2A deletions as we observed in CDKN2A^high^ tumours except most centered around the G2M checkpoint and transition instead of the G1-S checkpoint **(Figure 3g**,**i)**. Common down-regulated pathways included tumor necrosis factor (TNF), and cytochrome p450-related pathways **(Figure 3h**,**i)**.

### Downstream targets of the p16 pathway including CDK4 are aberrantly expressed in CDKN2A^high^ meningiomas

To investigate the effects of CDKN2A mRNA expression on its immediate downstream targets, we looked at the expression of cyclin-dependent kinase-4 (CDK4), CDK6, transcriptional factor E2F3, and retinoblastoma-1 (RB1) in each cohort. Although CDKN2A is normally a negative regulator of CDK4, there was significantly higher CDK4 mRNA expression in CDKN2A^high^ meningiomas compared to CDKN2A^low^ tumours across all 4 cohorts **(Figure 4a-d)**. There were no significant differences in the mRNA expression the other target genes save for CDK6 in the Bayley et al. cohort **(Figure 4b)**. In addition to CDK4, other E2F target genes CCND1, TP53, MCM2, and TK1 all showed a significant positive correlation with CDKN2A expression across all 4 cohorts **(Figure 4e)**.

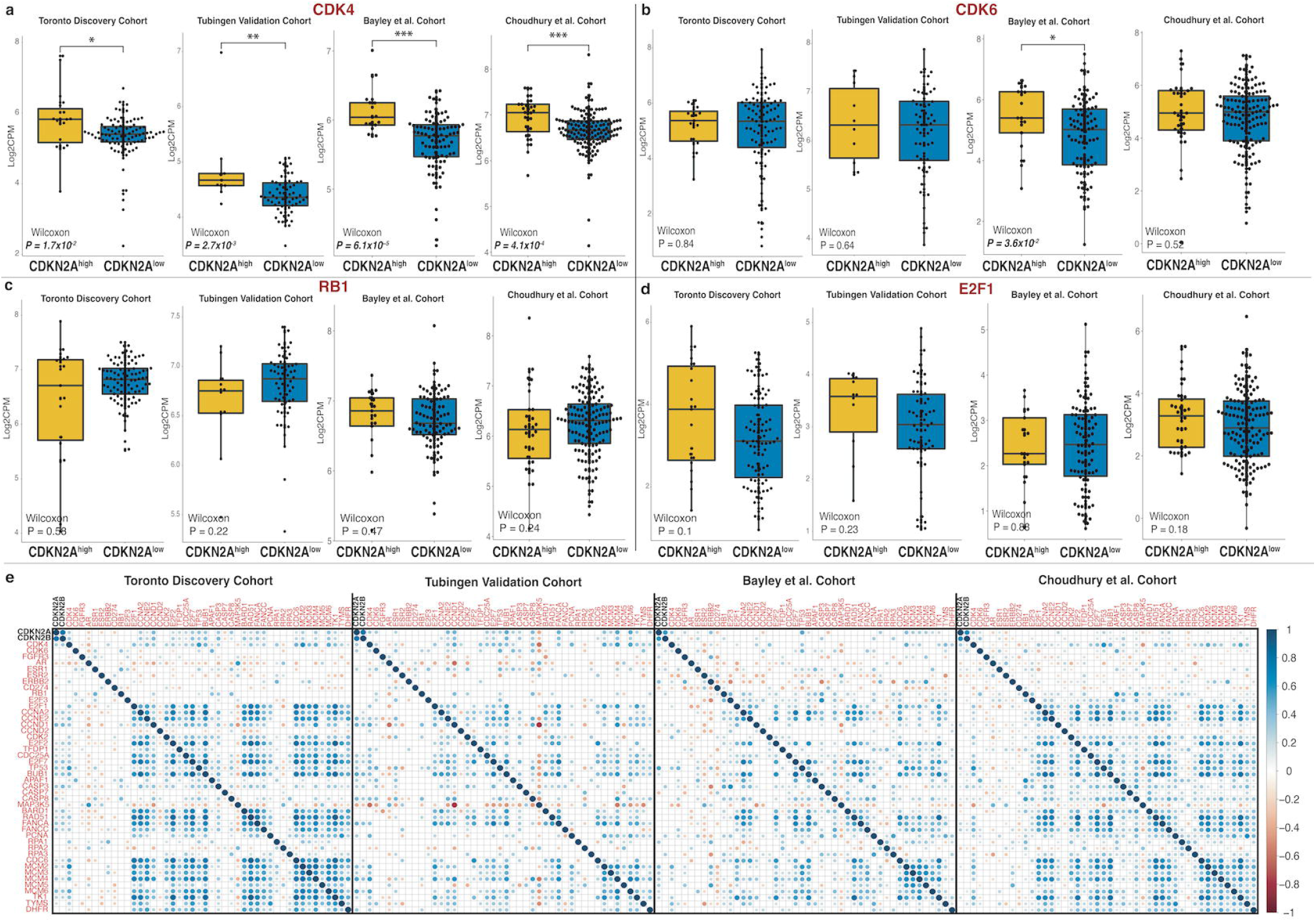

### Associated DNA methylation changes at the gene promoter and body of CDKN2A and CDK4

As CDKN2A and CDK4 are upstream of other E2F targets, and their expression appeared to be concordantly increased in CDKN2A^high^ meningiomas, we wanted to assess whether these transcriptomic changes could be correlated with DNA methylation at these loci. We found there was a significantly higher degree of methylation of CpG sites located at the CDKN2A gene, particularly at the gene body and 3’ untranslated region (UTR) in CDKN2A^high^ meningiomas compared to CDKN2A^low^ meningiomas across all 4 cohorts **(Figure 5a-d)**. These same methylation patterns were not seen for CDK4 **(Supplementary figure 4)**. On an individual CpG level, the methylation level of 14 different CpGs at the CDKN2A gene locus significantly correlated with CDKN2A mRNA expression in 3 or more cohorts **(Figure 5e-h**, p<0.05, Pearson’s correlation). Hypermethylation of four CpGs (cg08686553, cg14348664, cg16606671, cg26349275), located primarily in the 3’UTR and body of the CDKN2A gene, were significantly associated with increased CDKN2A expression in all 4 cohorts (**Figure 5e-h**, Pearson’s R = 0.43-0.94, p<0.05).

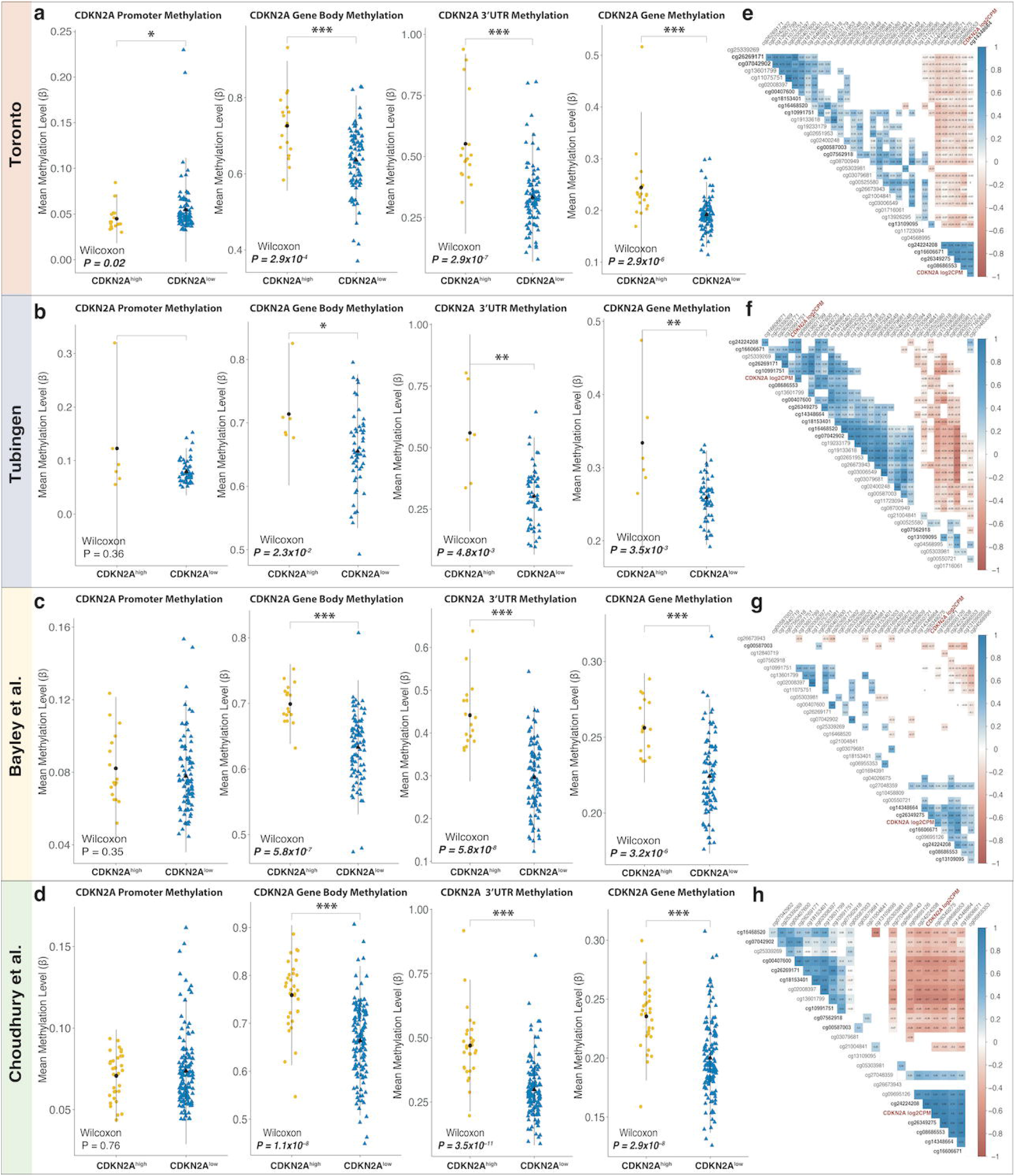

### Association of CDKN2A^high^ meningiomas with other copy number alterations

Next, we wanted to determine whether high mRNA expression of CDKN2A could be related to copy number amplification at its gene locus. However, there were no CDKN2A^high^ meningiomas with copy number gain of 9p at the chromosomal arm level or at the CDKN2A gene level **(Supplementary table 3, Supplementary figure 5)**, suggesting copy number changes are unlikely to be driving this increased expression. Copy number alterations of CDK4/6 were also rare and were only seen in a minority of meningiomas in the Toronto and Tübingen cohorts. **(Supplementary figure 5)**. There were a higher proportion of meningiomas with prognostically relevant copy number changes in the CDKN2A^high^ group vs CDKN2A^low^ across all cohorts including loss of 1p, 4p/q, 6p/q, 10p/q, 18p/q, and 22q **(Supplementary table 3, Supplementary figure 5)**.

### p16, CDK4 protein abundance corroborates transcriptomic data

Lastly, as the final step in the central dogma, we wanted to determine whether changes in gene expression of CDKN2A and CDK4 translated to differences in protein abundance. Overall p16 levels (the gene product of CDKN2A) appeared to increase with more aggressive MG, with MG4 meningiomas having the highest abundance of p16 protein detected **(Supplementary Figure 6a)**, concordant with our previously shown CDKN2A mRNA expression levels **(Figure 2a)**. When meningiomas were stratified by WHO grade, p16 also appeared to increase with higher WHO grade, although this trend was not statistically significant **(Supplementary Figure 6b)**. CDKN2A^high^ meningiomas were found to have the highest levels of p16 protein while meningiomas with CDKN2A homodel had the lowest **(Supplementary Figure 6c)**. p16 protein levels also showed significant positive correlation with CDKN2A mRNA expression (**Supplementary Figure 6d**, Pearson R=0.44, p=1.9×10^−6^). When p16 IHC were performed on a representative set of meningiomas with CDKN2A homodel but various levels of CDKN2A mRNA expression, those with higher levels of mRNA expression appeared to have a greater degree of positivity (nuclear and cytoplasmic), while those with low or null expression levels were challenging to differentiate from one another **(Supplementary Figure 6k)**.

CDK4 protein levels appeared to significantly increase with MG, in a similar manner to p16 **(Supp Fig 6e)** and were also significantly higher in WHO grade 2 meningiomas compared to WHO grade 1 tumors **(Supplementary Figure 6f)**. Similar to p16, CDKN2A^high^ meningiomas had the highest abundance of CDK4 protein compared to other groups **(Supplementary Figure 6g)**. However, unlike at the transcriptomics level, where meningiomas with CDKN2A homozygous deletion had the highest CDK4 mRNA expression, these meningiomas showed lower levels of detected CDK4 protein, although only 3 tumours in this group were profiled here. CDK4 protein abundance was also significantly and positively correlated with CDKN2A mRNA expression **(Supplementary Figure 6h**, Pearson R=0.42, p=5.8×10^−5^).

### Rb-deficiency may be more common in CDKN2A^high^ meningiomas

As phosphorylation of Rb (as opposed to its RNA expression or protein abundance) is a key nidus of control for cell cycle progression from G1 to S, we wanted to assess its phosphorylation status in representative meningiomas from each CDKN2A group. The Rb protein was present in all meningiomas with CDKN2A homodel (N=4) or heterodel (N=3) and were hyperphosphorylated at both key serine sites in all tumors (S780, S807/811, **Supplementary Figure 6i**). In CDKN2A^low^ meningiomas, Rb was present in all samples, but was only phosphorylated at both sites in 3/17 (17%) tumors. In CDKN2A^high^ meningiomas however, 58% of samples (N=7/12) were Rb-deficient and in 3/5 (60%) samples in which Rb was present, it was hyperphosphorylated at S780 and S807/811. This suggests that within CDKN2A^high^ meningiomas, there may be a higher proportion of Rb-deficient meningiomas and meningiomas with hyperphosphorylation of Rb that may behave similar to those with copy number loss of CDKN2A **(Supplementary Figure 6j)**.

## Discussion

CDKN2A is a gene located on chromosome 9 that encodes for two proteins: p16 and p14arf.^39,40^ Both these proteins act as tumor suppressors through regulation of the cell cycle. p16 normally inhibits CDK4 and CDK6, activating RB which blocks the G1-to S-phase transition while p14arf activates the p53 tumor suppressor.^39,41,42^ Therefore, loss of CDKN2A function or decreased expression in neoplastic conditions, should lead to unchecked cell-cycle progression, increased cell proliferation, and tumor progression.^11,18,43^ However, the fact that increased expression of CDKN2A has also been associated with poorer clinical outcomes in some cancers suggests that this relationship may be more complex.^13,18,19^ Although CDKN2A copy number loss has been found to be a marker of aggressive meningiomas, its relationship with the expression of its gene products is less clear. Here we demonstrated that only homodel, but not partial/heterodel of CDKN2A leads to significant reduction of CDKN2A gene expression, despite both alterations leading to similarly poor clinical outcomes, suggesting potential haploinsufficiency of the CDKN2A gene product.^44^ We also found that increased CDKN2A mRNA expression appears to be paradoxically associated with poorer PFS in meningiomas as well as higher WHO grade, and more aggressive molecular group in 4 independent cohorts. In investigating this potential phenomenon, we found that 1) CDKN2A^high^ meningiomas share transcriptomic pathways with meningiomas that have CDKN2A copy number loss, 2) changes in CDKN2A expression are associated with concordant methylation differences at the CDKN2A gene, 3) differences in CDKN2A at the mRNA level are mirrored by the relative abundance of the p16 protein, and 4) there are a subset of CDKN2A^high^ meningiomas that are Rb-deficient or have constitutive activation of the E2F pathway through Rb phosphorylation.

Several studies have suggested explanations for the discordant findings of CDKN2A/p16 overexpression in both benign/pre-malignant lesions and malignant tumors.^45^ While malignant transformation may be associated with early loss of CDKN2A in its tumor suppressive role, in the context of a functioning CDKN2A gene, aberrant tumour cell proliferation may result in incrementally increasing, compensatory expression of CDKN2A/p16 in a futile effort to halt cell cycle progression.^46,47,48^ This may explain the progressive increase in CDKN2A mRNA expression we observed in more proliferative meningiomas belonging to higher WHO grades and more aggressive molecular groups across multiple, independent validation cohorts. This too, may explain similarities in the upregulated transcriptomic pathways shared between meningiomas with CDKN2A deletion and CDKN2A^high^ meningiomas. While meningiomas with CDKN2A deletion and p16 bypass the G1/S checkpoint, progressing directly to G2 and mitosis due to constitutive activation of E2F, those with an intact CDKN2A gene must evade the G1 to S checkpoint and p53-mediated apoptotic pathways before cell cycle progression. The precise mechanisms by which this occurs remains uncertain, but may involve either loss of Rb, hyperphosphorylation of Rb, downstream alterations in the Rb pathway involving CDK4/CDK6 or CDK1/CCNA2, and/or Rb-independent pathways.

Interestingly, although CDK4/6 inhibitors have been proposed for aggressive, otherwise treatment-refractory meningiomas, their use has been suggested to be optimal for p16-deficient meningiomas (CDKN2A homodel) with an intact Rb.^49^ However, one of the limitations in using CDKN2A homodel as a prognostic marker is that this alteration is rare, and found in less than 5% of meningiomas, thereby failing to account for 15-20% of meningiomas that do not have this alteration, but may be similarly aggressive biologically. For these tumours, we propose that CDKN2A mRNA expression may be an effective biomarker that can be predictive of outcome independent of copy number loss and WHO grade. Additionally, though Rb-deficiency is rare in meningiomas, it may be more common in CDKN2A^high^ meningiomas, precluding the use of CDK4/6 inhibitors for these patients.^49,50^

By correlating our CDKN2A transcriptomic data with DNA methylation data, we found that increased methylation at the gene body and 3’UTR of the CDKN2A gene were associated with increased CDKN2A expression, particularly at several CpG sites that were consistently hypermethylated across all cohorts in CDKN2A^high^ meningiomas. Whether this represents a true regulatory mechanism for CDKN2A expression or whether these are simply passenger events in the context of known hypermethylation in more aggressive meningiomas, is uncertain. Multiple pan-cancer studies have shown however that gene body methylation and 3’UTR methylation are both associated with increased gene expression, and these may represent potential therapeutic targets for DNA methylation inhibitors.^51,52^

As RNAseq may be challenging to perform in a clinical setting, we showed a correlation between CDKN2A expression and p16 protein abundance that may be leveraged using IHC. A previous IHC study on 130 meningiomas found that increased p16 staining was strongly associated with poorer PFS on univariate analysis and that the proportion of meningiomas with p16 overexpression increased with increasing WHO grade, a seemingly concordant finding to our study.^53^ However, while p16 may not be optimal in differentiating meningiomas with CDKN2A homodel from those without any deletions, its positivity may correlate with CDKN2A mRNA expression. However, its utility needs to be confirmed in a larger cohort of meningiomas without CDKN2A deletion, and with matched RNAseq data.

Though we split each of our cohorts into CDKN2A^high^ and CDKN2A^low^ meningiomas, a universal threshold of CDKN2A expression that confers aggressive behaviour requires further investigation and validation across other large, independent cohorts. If, however, we treat meningiomas with CDKN2A homodel as having the lowest mRNA expression of CDKN2A, and CDKN2A^high^ meningiomas as those with the highest expression, then clearly CDKN2A^low^ meningiomas, that have no copy number loss of CDKN2A, and a relative intermediate level of CDKN2A mRNA expression, appear to have the optimal outcomes. However, the effect of partial or heterodel of CDKN2A on mRNA expression still requires investigation, ideally through methodology that compares next generation sequencing of a large cohort of meningiomas with array based CNV calling, as these tumours showed a highly variable degree of CDKN2A mRNA expression, despite their almost universally poor clinical outcomes. Lastly, functional studies that can assess changes in CDKN2A expression of meningioma cells with mitogenic stimulation may provide further mechanistic insights on the role of the CDKN2A-Cyclin-RB pathway in aggressive meningioma development and proliferation.

## Notes

### Competing Interest Statement

The authors have declared no competing interest.

