## Supplementary Materials for "Increased mRNA expression of CDKN2A is a transcriptomic marker of clinically aggressive meningiomas"

**Supplementary Table 1.** Summary of available matched clinical survival (progression-free survival; PFS) and molecular data (DNA methylation, RNAseq, proteomics) on the same patients from included cohorts and total.

| | **Toronto** | **Tubingen** | **Bayley et al.** | **Choudhury et al.** | ***Total*** |
|---|---|---|---|---|---|
| **DNA methylation** | 122 | 60 | 109 | 185 | ***476*** |
| **RNAseq** | 121 | 75 | 109 | 185 | ***490*** |
| **Protein** | 96 | 0 | 0 | 0 | ***96*** |
| **PFS data** | 122 | 75 | 109 | 160 | ***466*** |

**Supplementary Table 2.** CDKN2A expression group distribution across molecular groups in each cohort including only samples with RNAseq data.

| ***Toronto Discovery Cohort (n=121)*** | | | | | ***Tubingen Validation Cohort (n=75)*** | | | |
|---|---|---|---|---|---|---|---|---|
| | **Homodel (n=5)** | **Heterodel (n=6)** | **CDKN2A$^{low}$ (n=91)** | **CDKN2A$^{high}$ (n=19)** | **Homodel (n=0)** | **Partialdel (n=1)** | **CDKN2A$^{low}$ (n=64)** | **CDKN2A$^{high}$ (n=10)** |
| ***MG1*** | 0 | 0 | 14 (15%) | 3 (16%) | 0 | 0 | 5 (8%) | 0 |
| ***MG2*** | 1 (20%) | 3 (50%) | 28 (30%) | 0 | 0 | 0 | 25 (39%) | 1 (10%) |
| ***MG3*** | 1 (20%) | 1 (17%) | 36 (40%) | 5 (26%) | 0 | 0 | 25 (39%) | 2 (20%) |
| ***MG4*** | 3 (60%) | 2 (33%) | 13 (15%) | 11 (58%) | 0 | 1 (100%) | 9 (14%) | 7 (70%) |

| ***Bayley et al. Cohort (n=109)*** | | | | | ***Choudhury et al. Cohort (n=185)*** | | | | |
|---|---|---|---|---|---|---|---|---|---|
| | **Homodel (n=1)** | **Partialdel (n=1)** | **CDKN2A$^{low}$ (n=93)** | **CDKN2A$^{high}$ (n=14)** | | **Homodel (n=11)** | **Partialdel (n=5)** | **CDKN2A$^{low}$ (n=137)** | **CDKN2A$^{high}$ (n=32)** |
| ***MenG A*** | 0 | 0 | 47 (50%) | 4 (29%) | ***MI*** | 0 | 1 (20%) | 63 (46%) | 8 (25%) |
| ***MenG B*** | 0 | 0 | 23 (25%) | 3 (21%) | ***IE*** | 5 (45%) | 1 (20%) | 47 (34%) | 7 (22%) |
| ***MenG C*** | 1 (100%) | 1 (100%) | 23 (25%) | 7 (50%) | ***HM*** | 6 (55%) | 3 (60%) | 27 (20%) | 17 (53%) |

*HomoDel- homozygous deletion; HeteroDel- heterozygous deletion; MG- molecular group; MenG- meningioma groups; MI- Merlin-intact; IE- immune-enriched; HM- hypermitotic*

**Supplementary Table 3.** Frequency of common prognostic copy number alterations in CDKN2A$^{high}$ vs CDKN2A$^{low}$ meningiomas in each cohort

| | **Toronto** | | | **Tubingen** | | | **Bayley et al.** | | | **Choudhury et al.** | | |
|---|---|---|---|---|---|---|---|---|---|---|---|---|
| | CDKN2A$^{high}$ (n=19) | CDKN2A$^{low}$ (n=91) | P-val | CDKN2A$^{high}$ (n=6) | CDKN2A$^{low}$ (n=54) | P-val | CDKN2A$^{high}$ (n=17) | CDKN2A$^{low}$ (n=90) | P-Val | CDKN2A$^{high}$ (n=32) | CDKN2A$^{low}$ (n=137) | P-Val |
| ***1p-*** | 12 (63%) | 35 (38%) | 0.084 | 4 (67%) | 25 (46%) | 0.605 | 11 (67%) | 41 (46%) | **0.003** | 18 (56%) | 35 (26%) | **0.001** |
| ***4p-*** | 5 (26%) | 3 (3%) | **0.002** | 0 | 6 (11%) | 0.886 | 1 (6%) | 2 (2%) | 1 | 4 (13%) | 2 (1%) | **0.012** |

| | | | | | | | | | | | | |
|---|---|---|---|---|---|---|---|---|---|---|---|---|
| ***4q-*** | 3 (16%) | 1 (1%) | **0.014** | 0 | 5 (9%) | 1 | 1 (6%) | 1 (1%) | 1 | 3 (9%) | 4 (3%) | 0.247 |
| ***6p-*** | 5 (26%) | 7 (8%) | **0.049** | 1 (17%) | 4 (7%) | 1 | 0 | 2 (2%) | 1 | 5 (16%) | 5 (4%) | **0.030** |
| ***6q-*** | 10 (53%) | 14 (15%) | **0.001** | 1 (17%) | 7 (13%) | 1 | 3 (18%) | 6 (7%) | 0.308 | 8 (25%) | 17 (12%) | 0.126 |
| ***10p-*** | 5 (26%) | 5 (5%) | **0.015** | 1 (17%) | 3 (6%) | 1 | 0 | 1 (1%) | 1 | 1 (3%) | 2 (1%) | 1 |
| ***10q-*** | 6 (32%) | 9 (10%) | **0.033** | 1 (17%) | 4 (7%) | 1 | 0 | 2 (2%) | 1 | 3 (9%) | 3 (2%) | 0.148 |
| ***14q-*** | 6 (32%) | 19 (21%) | 0.477 | 2 (33%) | 14 (26%) | 1 | 3 (18%) | 3 (3%) | 0.075 | 6 (19%) | 21 (15%) | 0.836 |
| ***18p-*** | 5 (26%) | 11 (12%) | 0.214 | 1 (17%) | 4 (7%) | 1 | 3 (18%) | 1 (1%) | **0.009** | 3 (9%) | 14 (10%) | 1 |
| ***18q-*** | 7 (37%) | 13 (14%) | **0.046** | 1 (17%) | 5 (9%) | 1 | 5 (29%) | 2 (2%) | **0.0002** | 3 (9%) | 16 (12%) | 0.952 |
| ***22q-*** | 13 (68%) | 44 (48%) | 0.1802 | 4 (67%) | 39 (72%) | 1 | 12 (71%) | 28 (31%) | **0.005** | 21 (66%) | 58 (42%) | **0.029** |

*P-value from 2-sample test for equality of proportions with continuity correction.*

**Supplementary Figure 1.** PFS based on meningiomas stratified by CDKN2A/B copy number status in **a,** the Toronto discovery cohort (N=122) and **b,** the combined validation cohorts (Tubingen, Bayley et al., Choudhury et al.) that had available survival data (N=344). **c-d**, Pearson correlation of CDKN2A expression with CDKN2B in the Toronto cohort and the combined validation cohorts respectively, for samples where RNAseq data were available. **e-f,** violin plot of CDKN2A expression levels based on CDKN2A/B copy number deletion status in the Toronto discovery cohort and combined validation cohorts respectively. Adj. P from Kruskal Wallis test and post-hoc Dunn multiple comparisons test. *P<0.05.

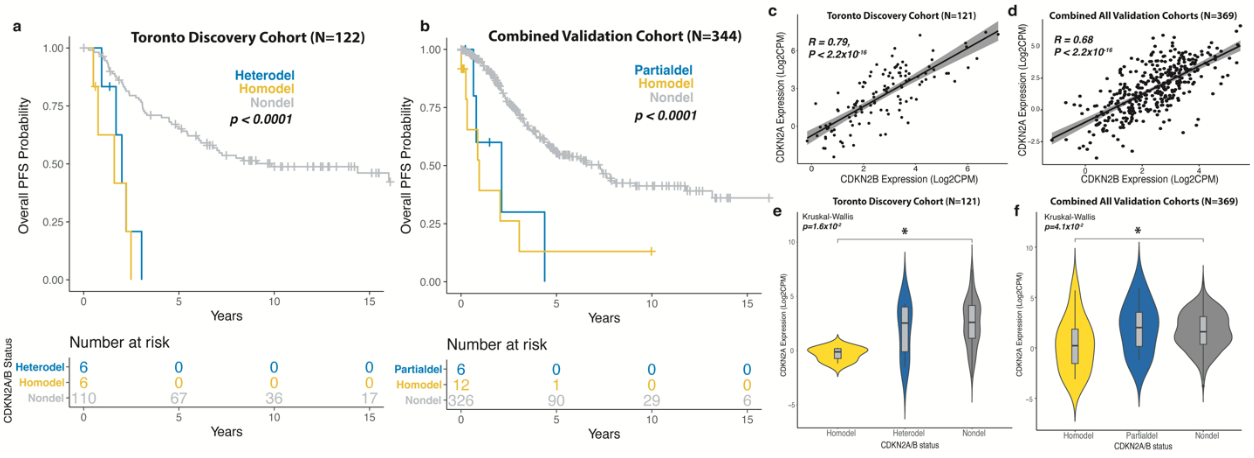


**Supplementary Figure 2.** Testing of the proportional hazards assumption for multivariable Cox proportional hazards model by plotting the scaled Schoenfeld residuals against transformed time for all covariates in the model.

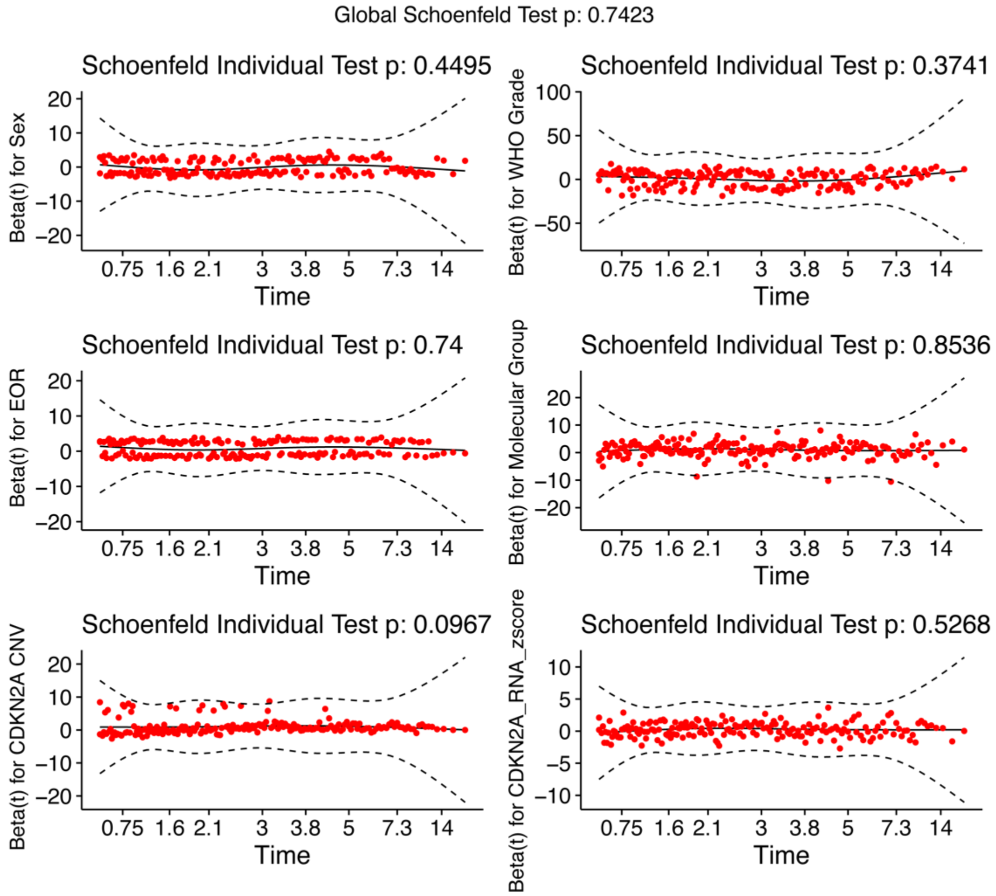


**Supplementary Figure 3. a-e,** mRNA expression counts in the combined cohort (Toronto, Tubingen, Bayley et al., Choudhury et al.) for E2F pathway genes based on CDKN2A status. Adj. P from Kruskal Wallis test and post-hoc Dunn multiple comparisons test. *P<0.05; **P<0.01; ***P<0.001.

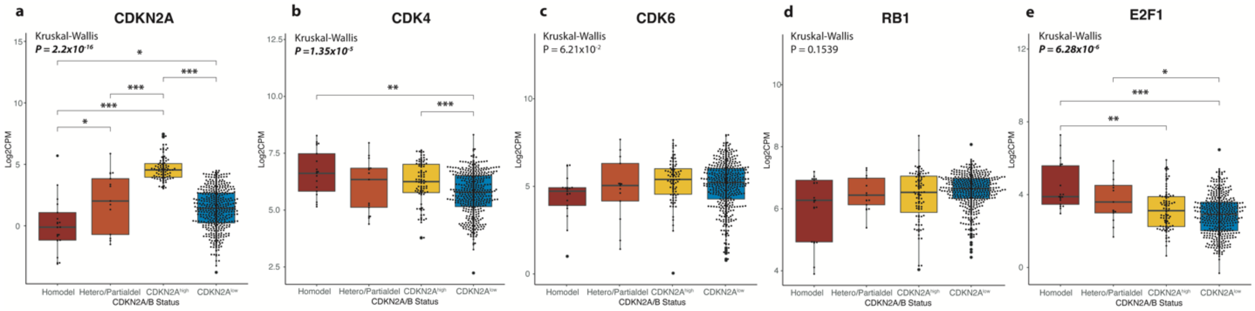

**Supplementary Figure 4.** Mean DNA methylation level at various regions of the CDK4 gene locus in A. the Toronto cohort, B. Tubingen cohort, C. Bayley et al. cohort, and D. Choudhury et al. cohort. Mann Whitney U-test *P<0.05.

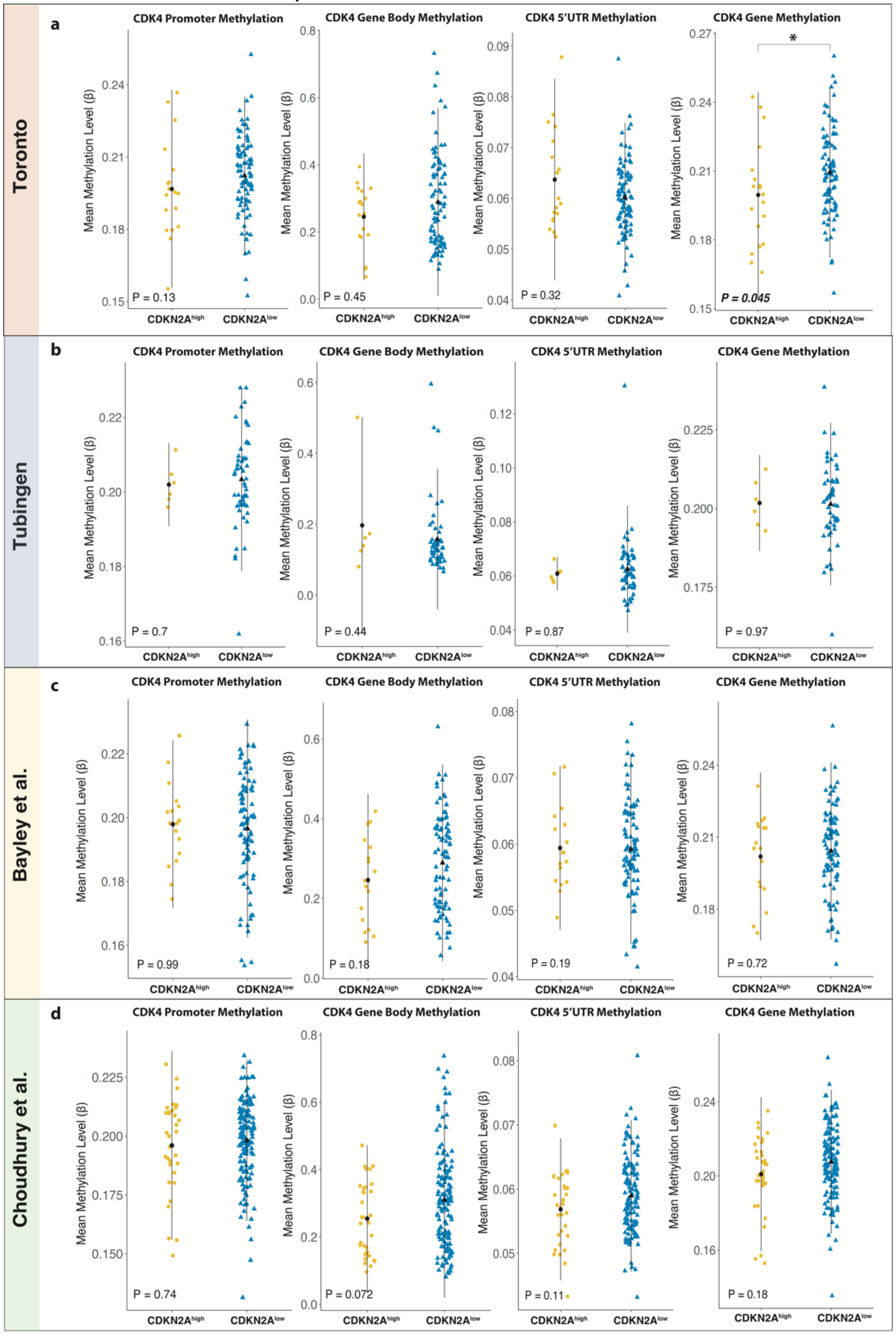

**Supplementary Figure 5.** Frequency of chromosomal arm-level and gene-level copy number alterations in each CDKN2A mRNA expression groups in the **A.** Toronto, **B.** Tubingen, **C.** Bayley et al., and **D.** Choudhury et al. cohorts.

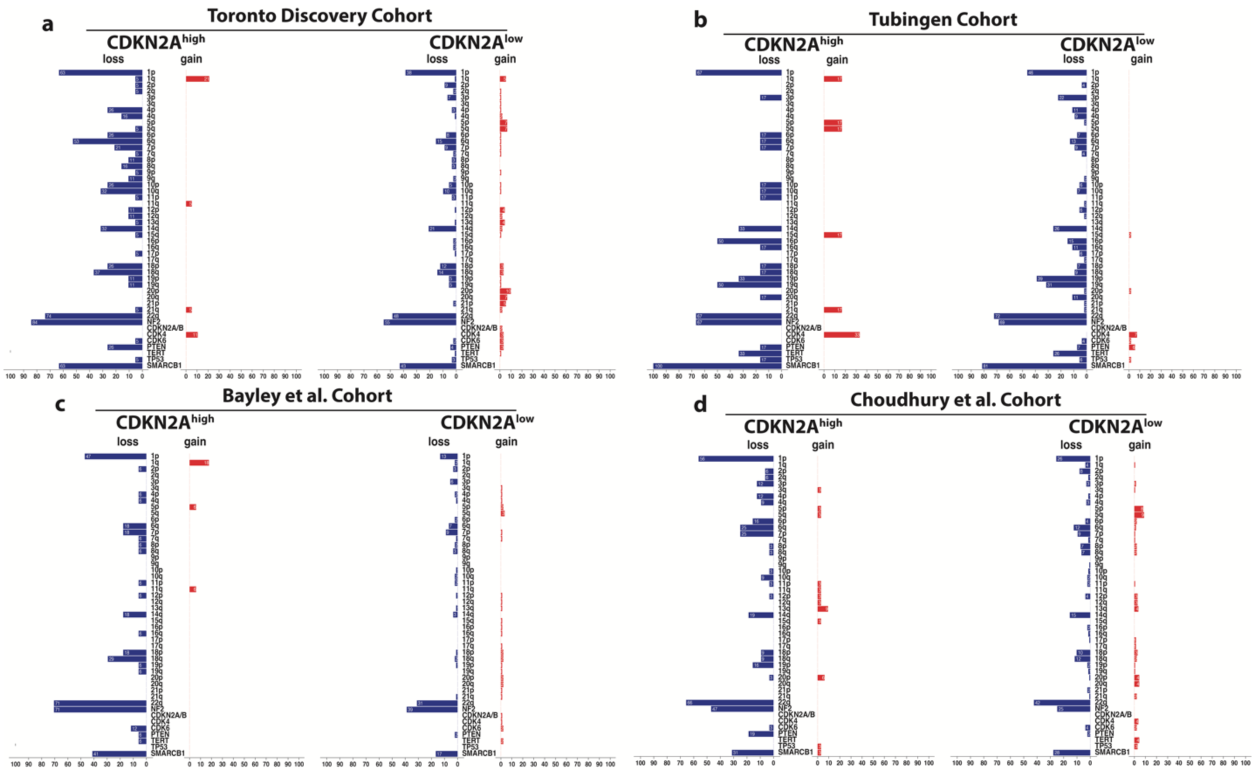

**Supplementary Figure 6**. Protein data from the Toronto cohort. Boxplot of p16 protein abundance with **a,** molecular group (MG), **b,** WHO grade, **c,** CDKN2A status (homozygous deletion, heterozygous deletion, $CDKN2A^{high}$, $CDKN2A^{low}$). **d,** Correlation plot of p16 protein levels vs CDKN2A protein levels. CDK4 Protein abundance by **e,** MG, **f,** WHO grade, **g,** CDKN2A status. **h,** correlation plot of CDK4 protein levels vs CDKN2A mRNA expression levels. **i,** Western blot of Rb phosphorylation at S780 and S807/811 in representative samples from each CDKN2A group. **j,** Effect of Rb phosphorylation or deficiency on cell-cycle progression. **k,** Representative p16 IHC from 6 meningiomas with CDKN2A homodel with different levels of CDKN2A mRNA expression

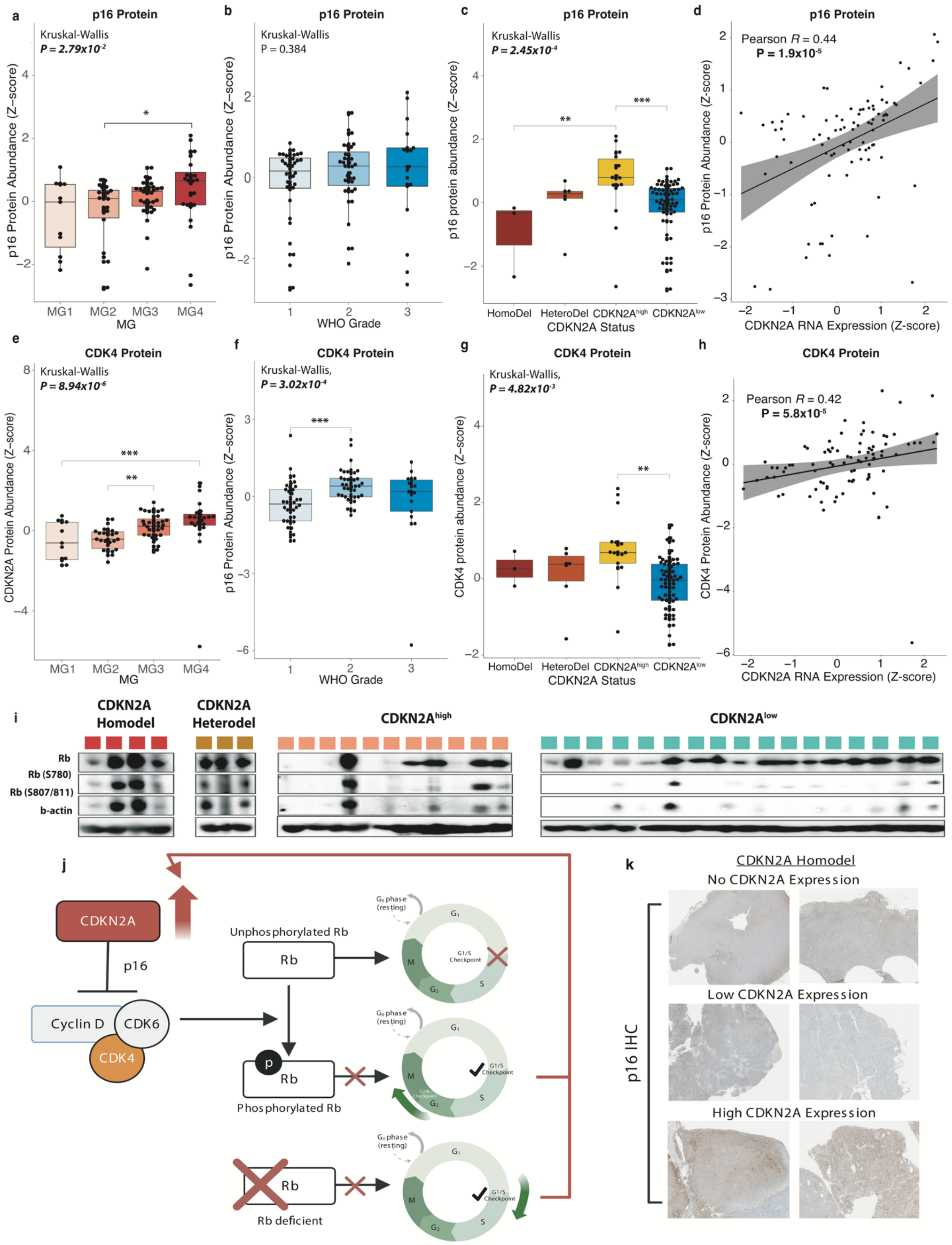

a
p16 Protein
Kruskal-Wallis
P = 2.79x10-2
p16 Protein Abundance (Z-score)
MG1 MG2 MG3 MG4
MG
b
p16 Protein
Kruskal-Wallis
P = 0.384
p16 Protein Abundance (Z-score)
WHO Grade
c
p16 Protein
Kruskal-Wallis
P = 2.45x10-4
p16 protein abundance (Z-score)
HomoDel HeteroDel CDKN2Ahigh CDKN2Alow
CDKN2A Status
d
p16 Protein
Pearson R = 0.44
P = 1.9x10-5
p16 Protein Abundance (Z-score)
CDKN2A RNA Expression (Z-score)
e
CDK4 Protein
Kruskal-Wallis
P = 8.94x10-6
CDKN2A Protein Abundance (Z-score)
MG1 MG2 MG3 MG4
MG
f
CDK4 Protein
Kruskal-Wallis,
P = 3.02x10-4
p16 Protein Abundance (Z-score)
WHO Grade
g
CDK4 Protein
Kruskal-Wallis,
P = 4.82x10-3
CDK4 protein abundance (Z-score)
HomoDel HeteroDel CDKN2Ahigh CDKN2Alow
CDKN2A Status
h
CDK4 Protein
Pearson R = 0.42
P = 5.8x10-5
CDK4 Protein Abundance (Z-score)
CDKN2A RNA Expression (Z-score)
i
CDKN2A Homodel
CDKN2A Heterodel
CDKN2Ahigh
CDKN2Alow
Rb
Rb (S780)
Rb (S807/811)
b-actin
j
CDKN2A
p16
Cyclin D
CDK6
CDK4
Unphosphorylated Rb
Rb
Phosphorylated Rb
Rb deficient
G0 phase (resting)
G1
G1/S Checkpoint
S
G2
M
k
CDKN2A Homodel
No CDKN2A Expression
Low CDKN2A Expression
High CDKN2A Expression
p16 IHC
